# Identification of a genomic DNA sequence that quantitatively modulates KLF1 expression in differentiating human hematopoietic cells

**DOI:** 10.1101/2021.11.22.469597

**Authors:** MN Gnanapragasam, A Planutis, JA Glassberg, JJ Bieker

## Abstract

Expression of the ß-like globin genes is under strict developmental control, with both direct and indirect inputs responsible for this effect. One of the major players regulating their transition is KLF1/EKLF, where even a two-fold difference in its level alters the regulation of globin switching. We have reproduced this change in KLF1 expression in both cell lines and primary human cells, thus demonstrating that directed, quantitative control of KLF1 expression can be attained by genomic manipulation, and suggest a new way in which modulation of transcription factor levels may be used for clinical benefit.

## Introduction

Hemoglobinopathies arise from deficient expression of globin chains (e.g., ß-thalassemia or Cooley’s anemia) or expression of mutant ß-globin (SCD). Together these are the most prevalent anemias caused by single gene mutations in the world (Taher et al., 2018). The associated morbidity is alleviated by expression of the dormant γ-globin (i.e., fetal) gene, which can follow from fortuitous genetic inheritance (Hereditary Persistence of Fetal Hemoglobin (HPFH)) or by its pharmacological induction with hydroxyurea (Kato et al., 2018; Ware et al., 2017). These alter the pattern of globin expression control during development, whereby switching of ß-like globin expression normally proceeds in an embryonic (ε-) to fetal (γ-) to adult (ß-) globin expression pattern.

Critical to the correct establishment of these changes are the controlling transcription factors, of which Erythroid Krüppel-like Factor (KLF1/KLF1) was the first to be discovered (Miller and Bieker, 1993). KLF1 accomplishes this feat by direct activation of the adult ß-globin and indirect repression of the fetal γ-globin gene (Bieker, 2010; Tallack and Perkins, 2013). KLF1 contains three C2H2 zinc fingers that precisely recognize its cognate DNA site (5’CCMCRCCCN3’) at target genes, enabling a high level of discrimination among similar sites (Donze et al., 1995). It performs this function by binding to its cognate DNA element, interacting with the basal transcription machinery, and recruiting chromatin remodeling proteins and histone modifiers (Siatecka and Bieker, 2011; Tallack and Perkins, 2010; Yien and Bieker, 2013). KLF1 integration of these signals leads to establishment of the correct 3-dimensional structure necessary for optimal target gene transcription (Drissen et al., 2004). Regulated expression of KLF1 target genes are coordinated within ∼40-60 nuclear transcription factories (Schoenfelder et al., 2010).

Although KLF1 interacts with corepressors (such as Sin3a and HDAC (Chen and Bieker, 2001)), the mechanism for its repression at the γ-globin promoter follows instead from its ability to activate proteins that directly repress γ-globin expression, such as BCL11A, ZBTB7A, SOX6, and KLF3 (Borg et al., 2010; Norton et al., 2017; Zhou et al., 2010).

Ablation of KLF1 leads to embryonic lethality in the mouse precisely at the time of the switch to adult ß-globin (Nuez et al., 1995; Perkins et al., 1995). This is only one of many changes that occur in the red cell in its absence, such that KLF1 is now recognized as a global regulator of all aspects of erythropoiesis (Perkins et al., 2016; Tallack et al., 2010). The range of phenotypic/clinical parameters that follow from its genetic modulation in the human has become extensively catalogued (Perkins et al., 2016; Waye and Eng, 2015). KLF1 haploinsufficiency leads to benign outcomes (e.g., lower expression of CD44 and Lu/BCAM antigen), but compound heterozygosity can lead to more serious problems such as pyruvate kinase deficiency/chronic nonspherocytic hemolytic anemia (Viprakasit et al., 2014) or to *hydrops fetalis* (Magor et al., 2015). Some mutations lead to a dominant phenotype such as seen in congenital dyserythropoietic anemia type IV (Arnaud et al., 2010; Varricchio et al., 2019).

Clinical parameters are altered when one KLF1 allele is inactive. Of particular interest to the present study, most prevalent is the loss of ß-like globin switching mechanisms, leading to HPFH (originally mapped in a Maltese family (Borg et al., 2010)) and also to persistence of embryonic globin (Viprakasit et al., 2014). There are also benign but significant changes in MCV and MCH (both lower), or in ZnPP or RTC (higher). These changes enable identification of KLF1 mutation carriers (Natiq et al., 2017; Perkins et al., 2016; Waye and Eng, 2015), and it is of considerable interest that allelic distribution of KLF1 mutations are dramatically higher in regions of endemic ß-thalassemia (Liu et al., 2014), where the dysregulation of γ-globin expression is of clinical benefit. A case report of ß°/ß° thalassemic twins highlights the remarkable clinical difference in anemia and transfusion dependence that follows from loss of one KLF1 allele (Xie et al., 2019). Collectively these data strongly suggest that directed modulation of KLF1 levels can provide a significant benefit to thalassemic and sickle cell patients.

KLF1 control elements have been clearly established by in vitro (Chen et al., 1998) and in vivo (Adelman et al., 2002; Xue et al., 2004; Zhou et al., 2010) experiments. In addition to its proximal promoter (Crossley et al., 1994), KLF1 is regulated by an upstream enhancer element that exhibits an erythroid-specific hypersensitive site and is bound by general and tissue-restricted regulators (Chen et al., 2019; Lohmann et al., 2015). Coupled to this, an intronic enhancer element plays a critical role in establishing its optimal expression level (Lohmann and Bieker, 2008). These studies have delineated a relatively simple layout of a short (∼1 kB) region adjacent to the transcriptional start site that contains erythroid DNAse hypersensitive sites (EHS1 and 2) and sequence elements that binds hematopoietic transcription factors and respond to BMP4 signaling.

We recently analyzed the KLF1 transcription unit for variability across of a range of leukemic and malignant erythroid subsets and found only common SNPs (Gnanapragasam et al., 2018). However, we had not examined Juvenile Myelomonocytic Leukemia (JMML) samples. Evaluation of the KLF1 transcription unit in JMML samples directed us to more precisely investigate the role of intron 1 in aiding the optimal level of expression of KLF1.

## Results

### Identification of a mutation at the intronic enhancer region in a JMML patient with increased HbF

We sought to investigate KLF1 gene status in cells from patients that exhibit high HbF. With this in mind, we turned to JMML, an early childhood myeloproliferative/myelodysplastic disease that is a clonal disorder of pluripotent stem cells (Emanuel, 2004; Stieglitz et al., 2015; Tefferi and Gilliland, 2007; Van Etten and Shannon, 2004). It has long been noted that many of these patients have abnormally high HbF levels (Papayannopoulou et al., 1991; Shimizu et al., 1988) that correlate with the course of the disease (Niemeyer and Flotho, 2019). They are also thrombocytopenic and exhibit hepatosplenomegaly (Emanuel, 2004). We sequenced the complete KLF1 transcription unit (Gnanapragasam et al., 2018) in genomic DNA from 20 JMML patient samples to search for mutations (Stieglitz et al., 2015). We identified three changes within the samples, two of which were in the upstream promoter (Table S1, Fig S1A). Of particular interest, one sample from a patient with a slightly higher than normal HbF (HM3554) contains a single heterozygous base mutation within a highly conserved region of intron 1 that is not a SNP (Table S1, Fig S1B). This change did not appear in our previous analysis of over 5000 genomes from direct sequencing and from public databases (Gnanapragasam et al., 2018). This mutation is located near the intronic enhancer region we had previously discovered in the mouse (Lohmann and Bieker, 2008). Inclusion of this intron leads to a 3-fold increase in linked reporter activity beyond that seen with the 5’-promoter alone, as assayed in differentiating mouse embryoid bodies derived from mouse embryonic stem cells (Fig 1A).

**Figure 1.**
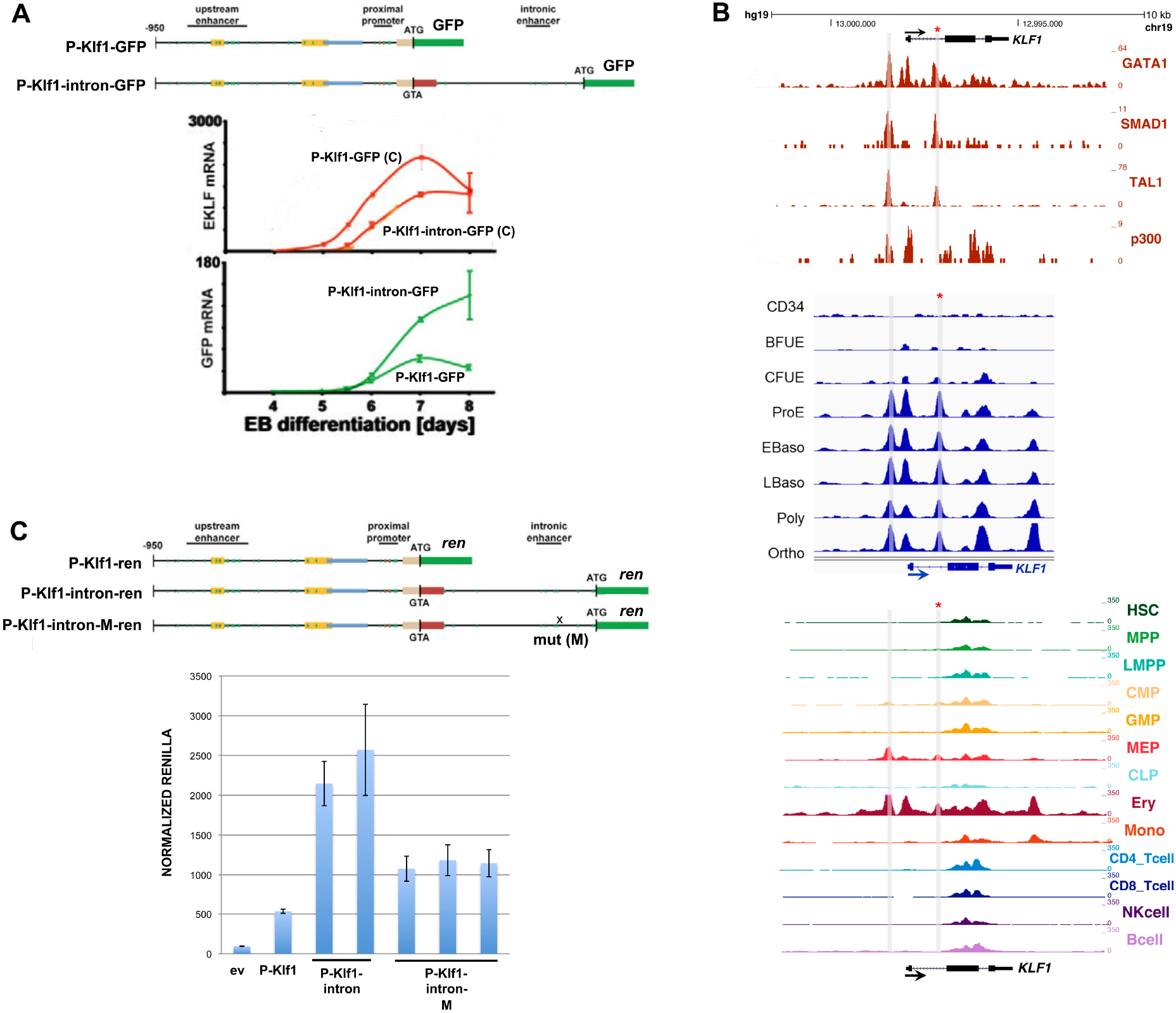
Analysis of the KLF1 intronic enhancer. (A) Reanalysis of data from a study (Lohmann and Bieker, 2008) on the mouse KLF1 promoter. Constructs containing the KLF1 promoter or one that additionally includes intron 1 were stably integrated into mouse ES cells adjacent to a GFP reporter, yielding two stable ES lines (“P-Klf1-GFP” and “P-Klf1--intron-GFP”). Top: Locations of the mapped upstream enhancer, proximal promoter, and intronic enhancer are as indicated. Bottom: ES cells from each line were differentiated to EBs for the indicated number of days, and samples were quantitatively analyzed for expression of endogenous KLF1 to monitor normal expression onset (red; C=control) or exogenous reporter (green) to monitor the effect of intron 1 inclusion. (B) Genome browser data aligned at the human KLF1 genomic region. Shaded bars show the locations of the enhancer upstream of the gene and of the intron 1 site within the gene (also marked with an asterisk). Top: In vivo GATA1, SMAD1, TAL1, and P300 binding identified from ChIP-seq analyses of erythroid cells. Middle: Onset of chromatin accessibility during erythroid differentiation from human CD34+ cells as monitored by ATAC-seq analyses of sorted cells (Schulz et al., 2019). Bottom: Comparison of chromatin accessibility (ATAC-seq) of prospectively isolated human hematopoietic populations (Ulirsch et al., 2019). (C) Reporter assay of various renilla reporter constructs after transfection into human JK1 cells. Top: Schematic of constructs containing the KLF1 promoter (as in (A)) are shown. The location of the intron 1 point mutant introduced into P-Klf1-intron-Ren is shown (“M”). Bottom: Results of the assay, showing high renilla levels from the promoter alone that are further stimulated by inclusion of the intron (two separate DNA preparations); however these levels are decreased when the point mutant variant is used (three separate preparations). Normalization is to a co-transfected luciferase plasmid, and data is an average of triplicate samples.

A closer analysis reveals that the mutation is located within a sequence that is highly conserved across mammalian species, and directly adjacent to previously identified GATA and SMAD recognition elements in the mouse (Lohmann and Bieker, 2008) (Fig S1B). This region binds GATA1, SMAD1, and TAL1 transcription factors also in human erythroid cells (Fig 1B), and the increase in chromatin accessibility during human hematopoiesis and particularly during erythroid differentiation as judged by ATAC seq analysis is centered at that site (along with that of the previously-described upstream promoter/enhancer) (Fig 1B) (Ludwig et al., 2019; Schulz et al., 2019; Ulirsch et al., 2019). These data suggest that the intron 1 site mutated in the JMML patient is within a functionally important element that is critical for optimal mammalian KLF1 expression.

### Reporter assays demonstrate the relevance of the intronic enhancer mutation on KLF1 expression

We performed reporter assays in transfected JK1 human erythroleukemia cell line to address the importance of the intron 1 mutation. This was assessed by transfection of renilla reporters driven by wild type or site-directed mutated KLF1 promoter and intronic regions (design based on (Lohmann and Bieker, 2008)). Fig 1C shows that, as expected from the murine data, inclusion of intron 1 increases the activity of the promoter ∼4-fold. However, this increase is reduced by 50% when performing the same assay using a reporter with the intron 1 single nucleotide mutation, verifying its importance. Performing the same assay with the two upstream promoter changes had little effect (Fig S2).

### Genome editing at the mutated intronic region in the KU812 human erythroleukemia cell line leads to reduced KLF1 and elevated epsilon globin expression

We next addressed whether the sequence surrounding the mutation is critical for hKLF1 transcription by using TALEN directed-nuclease technology (Joung and Sander, 2013; Miller et al., 2011) to delete the site within the KLF1 genome in human cells (Fig 2A), and monitored KLF1 levels. We were aided in the selection process by the use of a cotransfected RFP/GFP reporter (Kim et al., 2011) (Fig S3A). Using this system we transfected human KU812 cells which, although of low transfection efficiency via lipofection, enabled us to select for double positive RFP+/GFP+ cells (Fig S3A).

**Figure 2.**
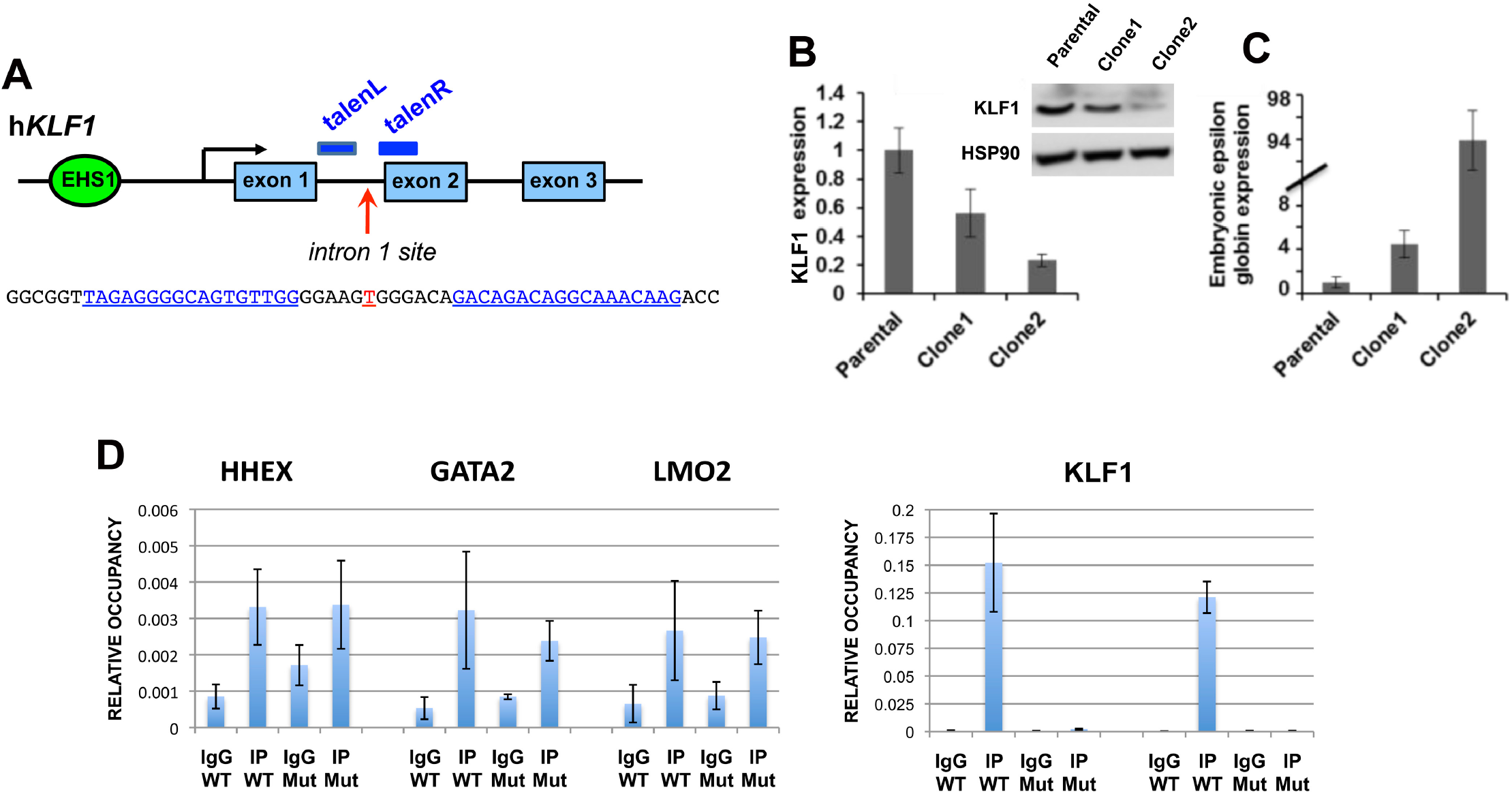
TALEN-mediated indel generation and analysis of KU812 cells. (A) Schematic of the location of the two TALEN arms (blue) in the context of the KLF1 gene and the intron 1 mutation site (red). Exact sequence is shown below. (B) Two of the KU812 cell clones (Fig S3B) compared to parental cells are shown after analysis of KLF1 RNA expression and western blot assessment of KLF1 and HSP90 protein (insert). (C) RNA expression of ε-globin was monitored in the KU812 clones [note scale change]. (D) ChIP analysis of specific TEL/ETV6 binding in WT or mutant (clone 2, “Mut”) KU812 cells. Left, positive control targets known to bind ETV6 show signals in all cases (HHEX, GATA2, LMO2) in WT or mutant cells. Right, ETV6 occupancy at the KLF1 intron 1 site, monitored by two different primer pairs, show a positive signal only WT cells. For all experiments, multiple samples were analyzed, each in triplicate.

The clones encompassed homo- and heterozygous deletions of 3 to 15 nt centered at the intron 1 mutation site (e.g., Fig S3B), indicating successful attainment of genomic editing. Clones of non talen-treated parental cells were also obtained to serve as a control to avoid any variables from clonal selection. Western and qRT-PCR analysis was performed for clone 1 (heterozygous deletion), clone 2 (homozygous deletion), and one of the parental clones (Fig S3B). Western analysis shows that KLF1 protein levels are quantitatively reduced in clone 1 and more reduced in clone 2 compared to a parental clone, effects that mirror the KLF1 transcript expression (Fig 2B). Globin expression analysis reveals that while ß- and γ-globin expression were unaffected (not shown), ε-globin levels are dramatically upregulated in the KU812 cell line in proportion to KLF1 expression (Fig 2C), consistent with the dysregulation of ε-globin seen in human KLF1 mutant patients with chronic nonspherocytic hemolytic anemia (Viprakasit et al., 2014).

Although γ-globin levels were not affected in this cell line (as they are already quite high to begin with in the parental cell (not shown)), these results enable us to sublocalize the functionally critical sequence from within the original large 0.9 kB intron 1 to a small ∼15 bp region surrounding the intron 1 site mutation, and suggest that a circumscribed deletion centered on this mutation site within normal cells will quantitatively decrease, but not ablate, KLF1 expression.

### Identification of TEL/ETV6 protein interaction with the KLF1 intron 1 region

Given the high level of sequence conservation at this site (Fig S1B), transcripton factor interactions may be predicted to be negatively affected by the mutation. We used the UniPROBE factor interaction prediction analysis to identify binding sites that are directly disrupted by the intronic mutation (Hume et al., 2015; Newburger and Bulyk, 2009) coupled to analysis via the NIH Roadmap Epigenomics Consortium (Fig S4). We find that the intron region surrounding the mutated site is enriched for binding sites for the ETS family of proteins (Hollenhorst et al., 2011; Wei et al., 2010) that interact with a consensus motif known to be important for the transition from stem to erythroid cells (Thoms et al., 2021). The single base intron 1 mutation is predicted to disrupt the binding for these proteins. Exclusion of those transcription factors not expressed in erythroid cells (analysis via ErythronDB (Kingsley et al., 2013) and BioGPS (Wu et al., 2016)) leaves only a small subset as possible contenders for positive binding such as ERG, ETV6, and ELF2. Of particular interest is TEL/ETV6, a protein that, similar to KLF1, has known roles in enhancing erythroid differentiation and hemoglobin synthesis (Eguchi-Ishimae et al., 2009; Noetzli et al., 2015; Songdej and Rao, 2017) and in inhibiting megakaryopoiesis (Bouilloux et al., 2008; Frontelo et al., 2007; Siatecka et al., 2007; Takahashi et al., 2005). In vivo ChIP analyses of WT and intron 1-edited KU812 cells show that TEL/ETV6 binds to this site, and that its binding is abrogated when the target site is mutated (Fig 2D). We conclude that the TEL/ETV6 protein joins GATA1/SMAD/TAL1 in forming an optimal protein/DNA complex at this critical site within intron 1 of the KLF1 gene.

### Genome editing of KLF1 intron 1 in human CD34+ cells

To test whether quantitative knock down of KLF1 expression could be attained by editing of intron 1 in primary cells, we expanded human CD34+ cells and transfected them with the verified TALEN arms, followed by differentiation towards the erythroid lineage using a three-step protocol (Antoniani et al., 2018; Breda et al., 2016; Grevet et al., 2018; Wu et al., 2019). Positively-transfected cells were selected and were cultured as a pool of cells. Sorted cells from transfections without the TALEN arms served as our unaltered control. Deep sequencing of the positive pool indicates that over 60% of the cells are edited at the expected deletion site that overlaps the intron 1 mutation (Fig 3A). Although encouraging, we found the resultant small deletions had little effect on KLF1 expression, possibly due to the fact that our efficiency was not near 100%, viability was not good after the DNA transfection, and at most we should have detected a 30% drop in expression (50% effect x ∼60% cells with deletion).

**Figure 3.**
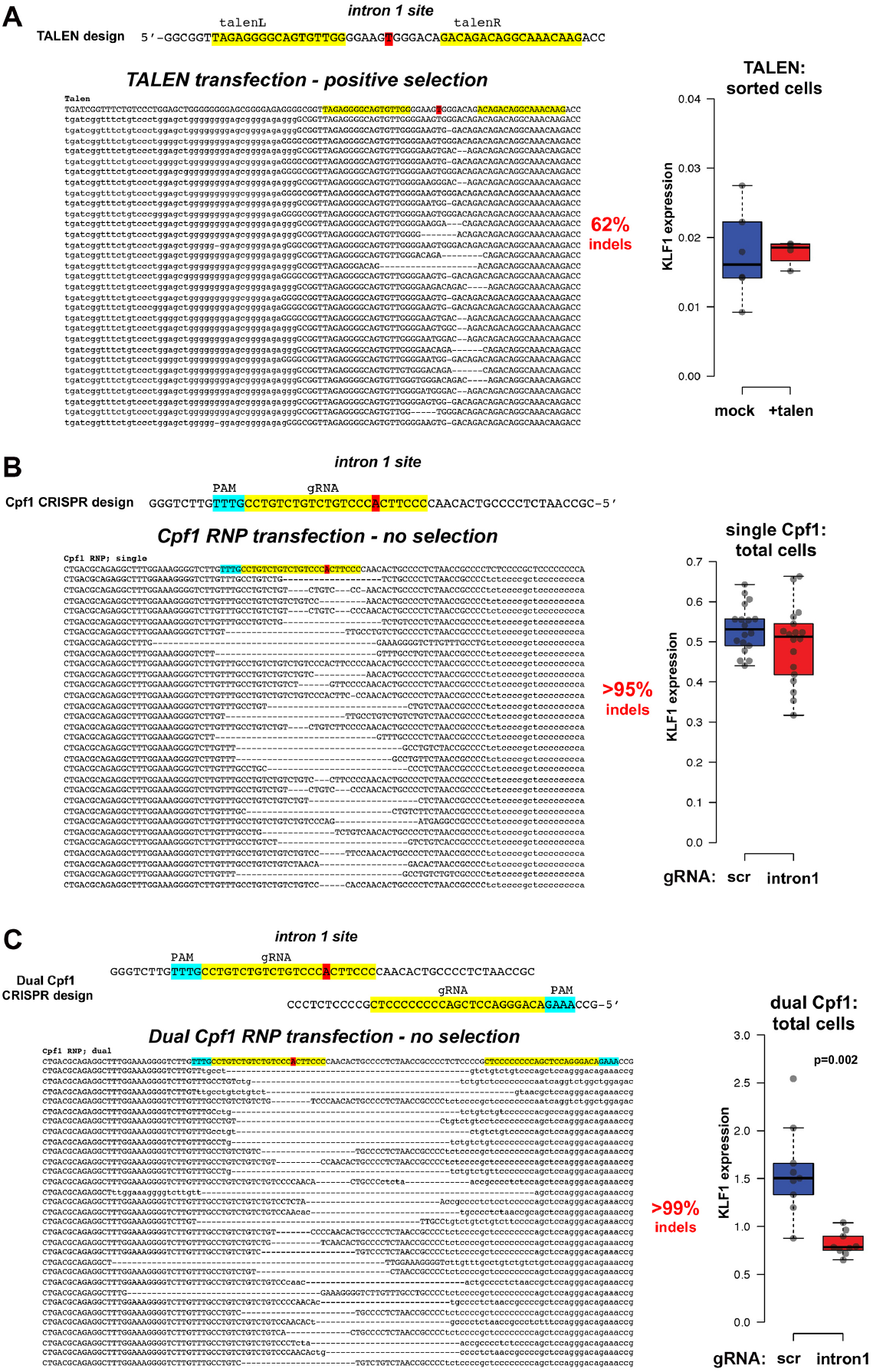
Gene editing and analysis of primary human cells. Human CD34+ cells were transfected, differentiated towards the erythroid lineage with a three-phase protocol, and analyzed by deep sequencing of the KLF1 intron 1 region. Indels were identified, and the top 30 are shown in each case along with the total indel percentage in the population. The location of the JMML point mutation (marked in red at the top) serves as a reference point for each design. KLF1 mRNA expression was monitored by RT-qPCR and shown on the right. (A) Cells were transfected with the left and right TALEN arms (indicated at the top in yellow) along with the co-transfected reporter. The RFP+/GFP+ pool was sorted, expanded, and differentiated prior to NGS genomic DNA analysis. KLF1 expression levels are not significantly different (n=4-6 each). (B) Cells were transfected with an RNP consisting of Cpf1 protein and gRNA (indicated at the top in yellow, with the PAM sequence in blue). The negative control was cells transfected with an RNP containing scrambled gRNA. The cells were expanded and differentiated prior to NGS genomic DNA analysis. NGS of the scrambled control cells showed no changes (not shown). Although there is a range of KLF1 expression after editing, levels are not significantly different from the scrambled control (n=18 each). (C) Cells were transfected with a 50/50 mix of RNPs consisting of Cpf1 protein and two gRNAs (indicated at the top in yellow, with their PAM sequences in blue). The negative control was cells transfected with an RNP containing scrambled gRNA. The cells were expanded and differentiated prior to NGS genomic DNA analysis. NGS of the scrambled control cells showed no changes (not shown). KLF1 expression is down 2-fold in the samples dual-edited samples (n=9 each).

During the initial design of these studies, we had not been able to use a Crispr approach due to the inherent requirements of Cas9 gRNA, whose PAM sequence (NGG) limited the target sequences to regions not near the site of interest in intron 1. However, in the interim Cpf1 (aka Cas12a) was discovered and developed as a viable alternate (Hur et al., 2016; Liu et al., 2020). Its merits in the present situation are that its PAM sequence (TTTV) is significantly different from that of Cas9, and that the *Acidaminococcus sp* variant has been engineered for increased activity and fidelity (Kleinstiver et al., 2019). This novel recognition sequence enabled us to design a Cpf1 gRNA that directly overlaps the site of interest and exhibits no predicted off-targets even with up to 2 mismatches (CRISPOR program; (Concordet and Haeussler, 2018)). We also opted to use an RNP-based approach to enable entry of the Cpf1/gRNA complex into CD34+ cells at high efficiency while retaining high cell viability (Gundry et al., 2016; Vakulskas et al., 2018; Weber et al., 2020; Wu et al., 2019). We tested this new design/approach, and indeed find that, even in the absence of selection, >95% of the cells are edited (Fig 3B). Use of a scrambled Cpf1/gRNA control routinely gives close to zero editing. The deletions at the region of interest are larger than had been observed with TALENs, but surprisingly we still do not see a significant effect on KLF1 levels on average even though the range of effect was wide (i.e., a subset of the data points were decreased compared to the control).

We next considered whether the Cpf1-directed deletion at the KLF1 intron 1 site might be too limited. Although one might predict that disruption of transcription factor binding should have a major effect on activity of enhancer function, it has been observed that removal of a single site in vivo can have a surprisingly minor effect (concepts discussed in (Blobel et al., 2021)), as any associated ‘enhanceosome’ components may still form if those protein-protein interactions are strong, and if there are other adjacent DNA binding sites that retain recruitment and cooperativity of other components in the full complex. Indeed, the KLF1 gene forms an extended 3D complex with multiple enhancers in its vicinity (Hua et al., 2021). We inspected the sequence surrounding the original Cpf1 position and found an additional Cpf1 PAM-sequence target site ∼35 bp away that also exhibits no predicted off-targets even with up to 2 mismatches (CRISPOR program; (Concordet and Haeussler, 2018)). We postulated that introduction of both Cpf1 gRNA sequences might enable a larger deletion region to be generated. This dual Cpf1 approach was tested. Again we attain high efficiency of editing in the absence of selection (>99%), but we now create a larger deletion and produce the anticipated 50% drop in KLF1 levels (Fig 3C). This suggests that focused disruption of the ETV6 site is not sufficient, but rather that deletion of a slightly larger region, perhaps containing other important binding sites, is needed to disrupt the enhancing activity of intron 1. Importantly, this verifies the importance of the intron 1 region for optimal KLF1 expression in vivo in primary cells and sets the stage for detailed analysis of its downstream effects.

## Discussion

We have identified a short, conserved enhancer element in KLF1 intron 1 that is important for establishing optimal levels of KLF1 expression in mouse and human cells. Further, we have developed a dual Cpf1 approach to editing this region that has extremely high efficiency and achieves exactly the drop in KLF1 levels that is likely to produce clinical benefit for hemoglobinopathies. Chromatin accessibility of this site exhibits cell-type specificity and is under developmental control during the differentiation of human CD34+ cells towards the erythroid lineage. This site binds GATA1, SMAD1, and TAL1, and we now show that it also binds ETV6. As ablation of KLF1 is lethal in mice and likely in humans, the identification of a site that quantitatively, rather than completely, decreases levels of KLF1 expression provides a novel target for gene therapy approaches designed to alleviate hemoglobinopathies.

The importance of quantitative control of expression is supported by the biology of KLF1 where too little creates problems, and by a recent study that depleted KLF1 in human CD34+ cells with two Crispr-directed targets located in exons 2 and 3 (Lamsfus-Calle et al., 2020). Although HbF was increased to similar levels as that seen after editing of the BCL11A enhancer or the HBG1/2 promoter, there were too many other genes dysregulated in their expression. This is not surprising given the known biology of KLF1. The present design is critically based on the observation that a haploinsufficient level of KLF1 is benign, and that select targets are dramatically affected by the 50% drop in expression (Siatecka and Bieker, 2011; Singleton et al., 2008; Singleton et al., 2012; Tallack and Perkins, 2013). Monoallelic disruption of KLF1 in humans is sufficient to knock-down the BCL11a repressor of γ-globin to the point where γ-globin is derepressed (Borg et al., 2010; Zhou et al., 2010). This was first mapped in a Maltese family that exhibited HPFH, and has since been observed in an increasingly large number of patients (reviewed in (Perkins et al., 2016; Waye and Eng, 2015)). An additional benefit to targeting KLF1 is that the ZBTB7A/LRF γ-globin repressor (Masuda et al., 2016) is another KLF1 target (Norton et al., 2017),

KLF1 expression is highly restricted to erythroid progenitors and progeny, as evidenced during development (Southwood et al., 1996), in the adult (Miller and Bieker, 1993), and during hematopoiesis (Cui et al., 2009; Frontelo et al., 2007; Li et al., 2012). The highly restricted nature of its expression is a crucial property, as quantitative alteration of its levels is predicted to limit its effect to the erythroid compartment, a unique situation that avoids the complexities that arise in other potential targets that are more generally expressed and required (Bauer et al., 2012; Brendel et al., 2016; Lazarus et al., 2018; Liu et al., 2003; Yu et al., 2012).

The inclusion of ETV6 into a positive regulatory role is perhaps a surprise, as it has been type cast as a transcriptional repressor. However as described before, its molecular role is complex, and in addition a recent study shows that KLF1 levels are decreased in homozygous mutant ETV6 iPS cells that were established as a model of inherited thrombocytopenia (Borst et al., 2021). These cells also exhibited decreased erythroid expansion capability and erythroid colony counts.

Our discovery that focused deletion of the ETV6 site was not sufficient to reliably decrease KLF1 levels, but that a larger deletion was required, makes the dual Cpf1 approach a defined way to achieve our aim of quantitatively knocking down KLF1 levels two-fold. Such a robust design may also avoid genetic modifier issues whereby there is an extensive level of variation seen in induction of HbF levels in patients containing the same monoallelic KLF1 mutation (e.g., (Eernstman et al., 2021)).

## Methods

KU812 and JK1 cells lines have been described (Drexler, 2010). Patient samples were as previously published (Stieglitz et al., 2015). Genomic sequencing using primers spanning the complete KLF1 transcription unit, including the upstream promoter, was performed in both directions as previously described (Gnanapragasam et al., 2018).

Reporter assays were performed after cotransfection of renilla reporter constructs into JK1 cells using X-tremegene HP transfection reagent (sigma) along with a luciferase expressing plasmid for normalization (Siatecka et al., 2007). Assays were performed using the dual luciferase kit (Promega) according to manufacturer’s instructions. Mutagenesis was performed with the Quick-change kit (Stratagene).

Mouse embryonic stem cell culture and differentiation into embryoid bodies was performed as described (Lohmann and Bieker, 2008) using cell lines generated by targeting the P-Klf1-GFP or P-Klf1-intron-GFP constructs into the site-specific Ainv18 homing site (Lohmann and Bieker, 2008). These contain a single copy, unidirectionally inserted sequence into the same site and avoids random integration.

RNA isolation and RT-qPCR analysis was as previously described (Gnanapragasam et al., 2016; Lohmann and Bieker, 2008).

CD34+ cells were purchased from AllCells or obtained from the Yale Cooperative Center of Excellence in Hematology. These were differentiated under three-phase protocols, initially based on (Antoniani et al., 2018; Yien and Bieker, 2012) but then later based on (Breda et al., 2016; Grevet et al., 2018; Wu et al., 2019). Transfection was performed after the expansion phase I, and cells were allowed to recover for 24-48 hrs in phase I media prior to switching to phase II.

Cloning of left and right TALEN arms was performed by insertion into a CMV promoter-containing plasmid, downstream of T7 polymerase promoter, HA tag, and NLS sequences, but upstream (in frame) of the Fok1 nuclease and bGH poly(A) signals (PNA Bio Inc). In addition, an RFP/GFP surrogate reporter was synthesized that contained the intron 1 sequence between them as a linker, but out of frame for GFP. Proper nuclease activity after transfection/expression leads to frameshift mutations and expression of GFP, enabling enrichment of a low number of positively transfected cells by sorting for RFP+/GFP+ double positivity (Fig S3A) (Kim et al., 2011).

Transfection of TALEN DNAs into KU812 cells was performed using the Neon Transfection System (1450v, 10ms, 3 pulses), sorted for RFP+/GFP+, and the positive pools were diluted to single cells into 96-well plates. After expansion, DNA was isolated and analyzed by PCR at the region of interest (using sequencing primer pairs (Gnanapragasam et al., 2018)) to determine whether any of these clones contained a deletion. Individual clones that were positive were expanded and used for subsequent analyses. Genomic PCR of cell line material used 25 ng purified DNA (Qiagen), followed by direct sequence analysis of the product (Macrogen).

CD34+ cells were transfected with the Amaxa Nucleofector II using program U-008 for TALEN DNA. For RNP we used the Amaxa 4D Nucleofector with program EO-100. Biological replicates were each analyzed in triplicate.

AsCpf1 protein (enhanced (Kleinstiver et al., 2019)) was purchased from IDT. RNP were freshly formed by mixing 105 pmol of Cpf1 protein with 120 pmol of gRNA (IDT) for 15 at room temperature, and kept on ice until use.

Positive pools of CD34+ cells were analyzed by next generation sequencing (MGH DNA Core Facility) following genomic DNA isolation and PCR. Alternatively, TIDE (Brinkman et al., 2014)or ICE (Hsiau et al., 2019) analysis was performed on these or on KU812 cell DNA samples using sequencing primer pairs (Gnanapragasam et al., 2018).

Predicted off-target effects of the TALEN pair was tested using the PROGNOS analysis program (Fine et al., 2014). We found no predicted off-targets when 0, or even 3, mismatches are allowed per arm (Table S2) (Fine et al., 2014). Predicted off-targets for the two gRNA designs used the CRISPOR program (Concordet and Haeussler, 2018), which showed no predicted off-targets even allowing for up to 2 mismatches (Table S3).

Chromatin immunoprecipitation of TEL/ETV6 was performed with a mix of antibodies (containing anti-TEL from Santa Cruz sc-1668335 and sc-8547 along with home-made rabbit polyclonal; generous gifts from Drs B Graves and K Clark (Hollenhorst et al., 2011)) using standard conditions. Positive control targets were chosen based on (Unnikrishnan et al., 2016).

We initially considered mutating the intron 1 region in HUDEP2 cells (Kurita et al., 2013), as they have been widely utilized for human erythroid studies. However, we obtained aberrant results with all our directed mutagenesis attempts at the KLF1 genomic region. Karyotype analysis revealed genomic instability along with trisomy 19 (not shown), which is the genomic location for KLF1. As a result, we switched to using JK1 and/or KU812 human leukemia cell lines for our initial set of gene editing analyses, as both of these also express KLF1 (unlike K562) and appear more stable (Bieker, 1996; Li et al., 2014).

## Acknowledgements

We thank Dr Mignon Loh (UCSF) for JMML patient samples. We thank Drs Barbara Graves and Kathleen Clark (University of Utah) for antibodies and discussion related to ETV6, and Dr John Pimanda (Prince of Wales Hospital) for target binding sequence information. We thank Drs Vesna Najfeld and Joseph Tripodi (MSSM) for karyotype analysis, and Drs Vivian Simon and Lubbertus Mulder (MSSM) for Amaxa usage. This work was supported by a Doris Duke Charitable Foundation award to JJB and JAG, by PHS grant R01 DK46865 to JJB, and by a Cooley’s Anemia Foundation Fellowship to MNG. The Yale Cooperative Center of Excellence in Hematology is supported by DK106829.

## Figure legends

**Figure S1.**
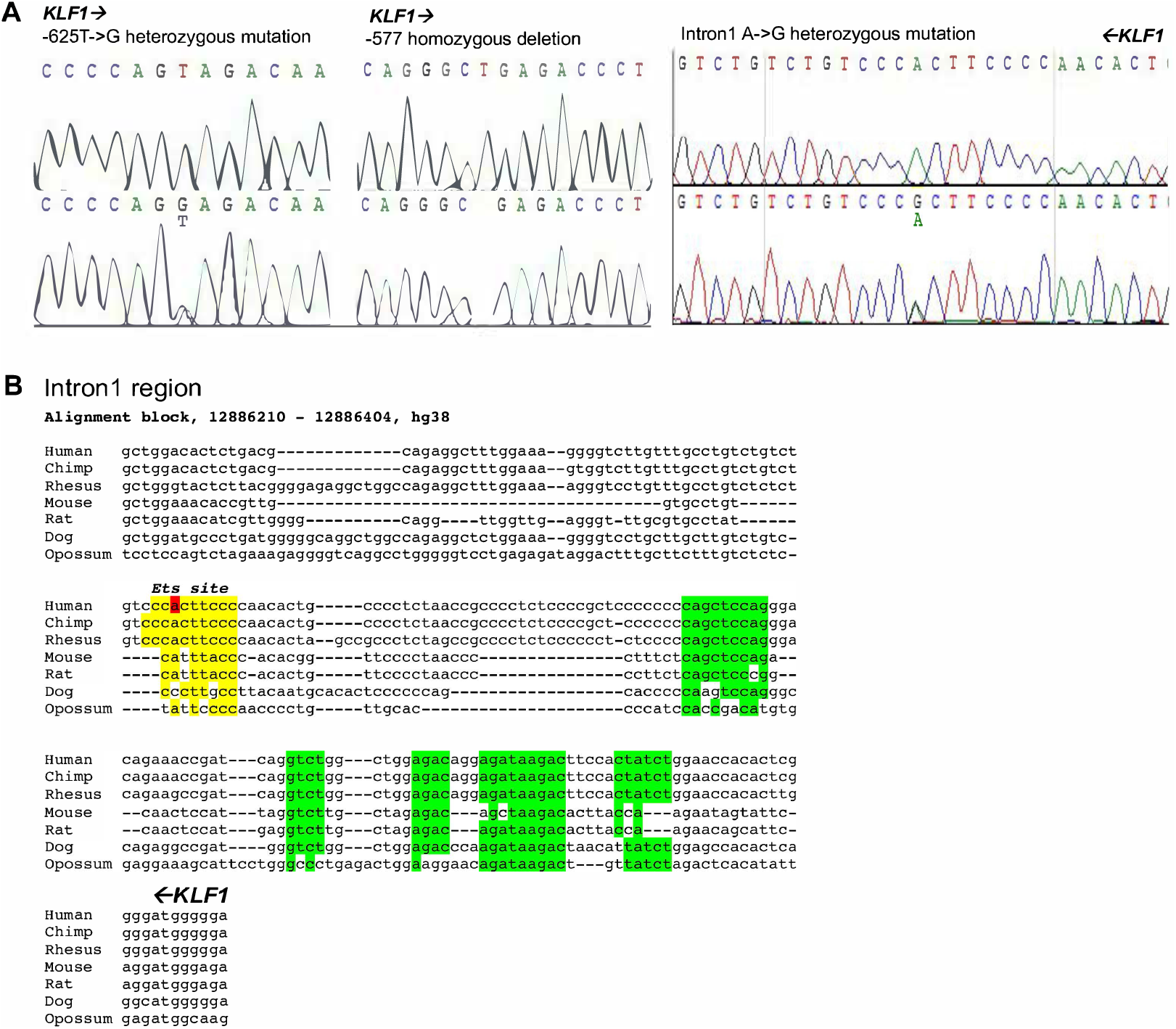
Sequence analysis of mutations identified from the JMML samples. (A) Sequence traces of regions surrounding −625, −577, and intron 1 mutation locations, showing WT sequence on top and mutant sequence below, with substitution/deletion as indicated. Orientation of KLF1 gene is indicated by direction of arrows. (B) Conservation of intron 1 region across selected vertebrates, with KLF1 gene orientation from right to left as indicated. The mutation is indicated in red, as is the putative ETS binding site in yellow. Green are other previously identified conserved regions (Lohmann and Bieker, 2008).

**Figure S2.**
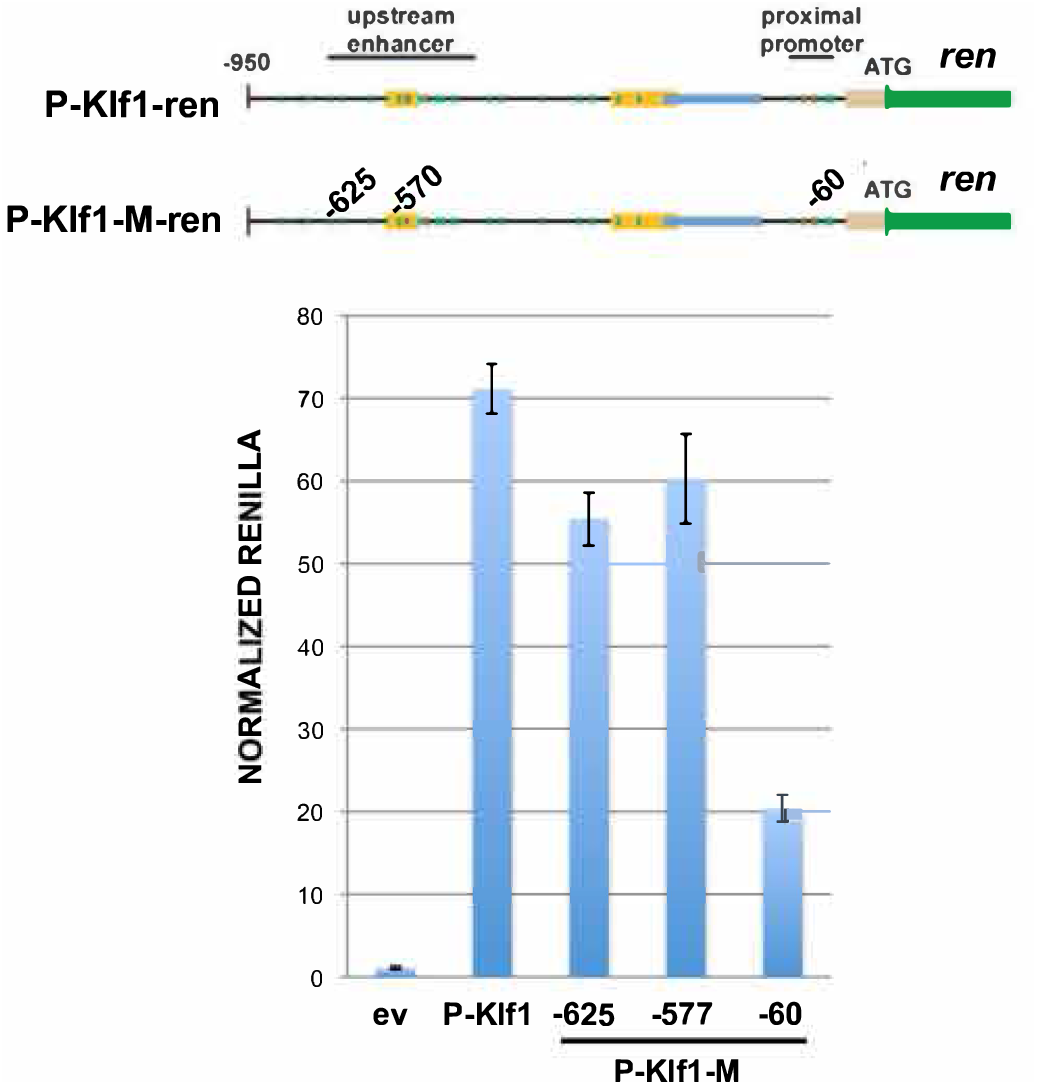
Reporter assay of various renilla reporter constructs after transfection into human JK1 cells. Top: Schematic of constructs showing the location of the JMML mutants introduced into P-Klf1-intron-Ren (“M”). Bottom: Assay results show that high renilla levels from the promoter are not significantly altered after inclusion of the −625 or −577 mutation; a reporter containing the −60 mutation (located at an important GATA site (Crossley et al., 1994)) is included as a positive control. Normalization is to a co-transfected luciferase plasmid, and data is an average of triplicate samples.

**Figure S3.**
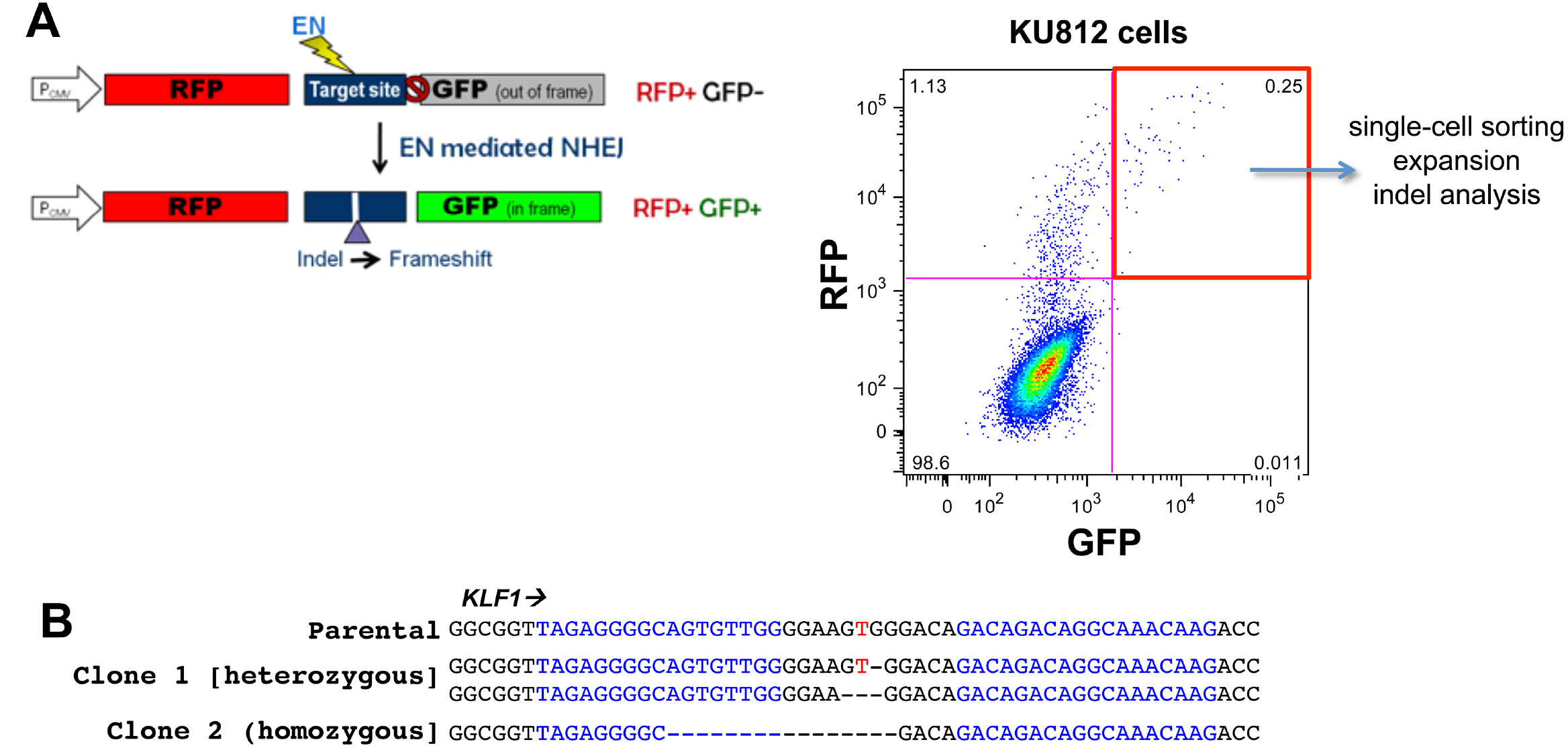
Strategy for isolating TALEN transfected/modified cells. (A) Left, schematic of parental (Kim et al., 2011) and modified constructs containing RFP (in frame) and GFP (out of frame) reporters separated by a target site into which the intron 1 sequence (with the left and right TALEN sites) is incorporated. Successful transfection and activity of the dual TALENs is predicted to yield an indel leading to a frameshift and resultant in-frame expression of GFP (1 out of 3 chance). Selection of RFP+/GFP+ positive cells after cotransfection with left and right TALEN arms greatly increases the chance of finding a clone with the desired indel, even at low frequency. Clones are generated by single-cell sorting and expansion. Data is shown from the transfection of KU812 cells. (B) Genomic sequence analysis of parental and two KU812 subclones containing directed indels at KLF1 intron 1; blue are the TALEN sequences, red is the target JMML site as a reference point. Homo- or heterozygous deletions are as indicated.

**Figure S4.**
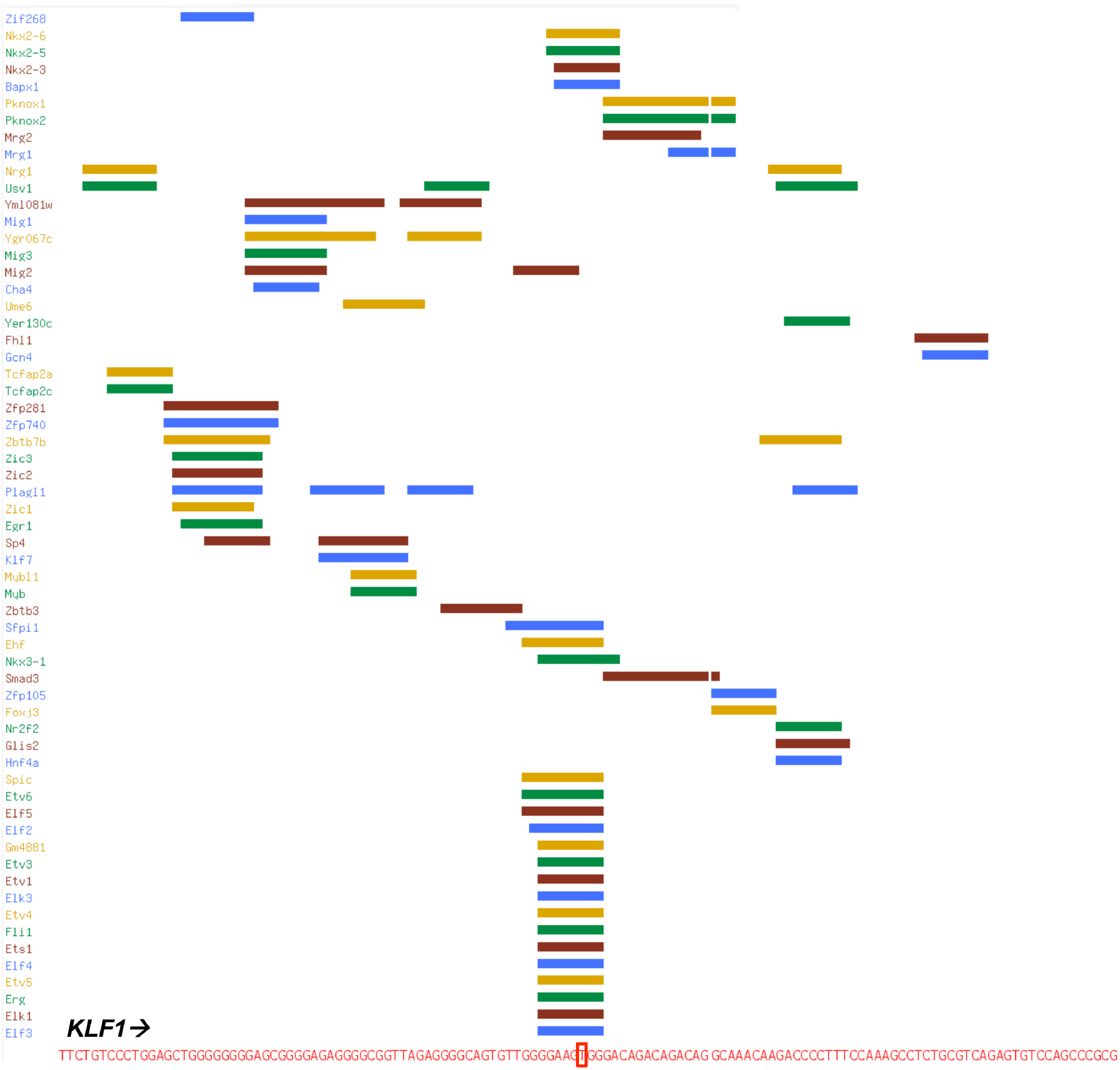
UniPROBE analysis of the region surrounding the intron 1 site. 100 nucleotide sequence surrounding the intron 1 site of interest (marked in red; oriented from left to right) was searched by the UniPROBE program (Hume et al., 2015; Newburger and Bulyk, 2009) to identify potential cognate binding transcription factors. Among the factors predicted to recognize the intron 1 site are multiple members of the Ets family of proteins.

**Table 1.**
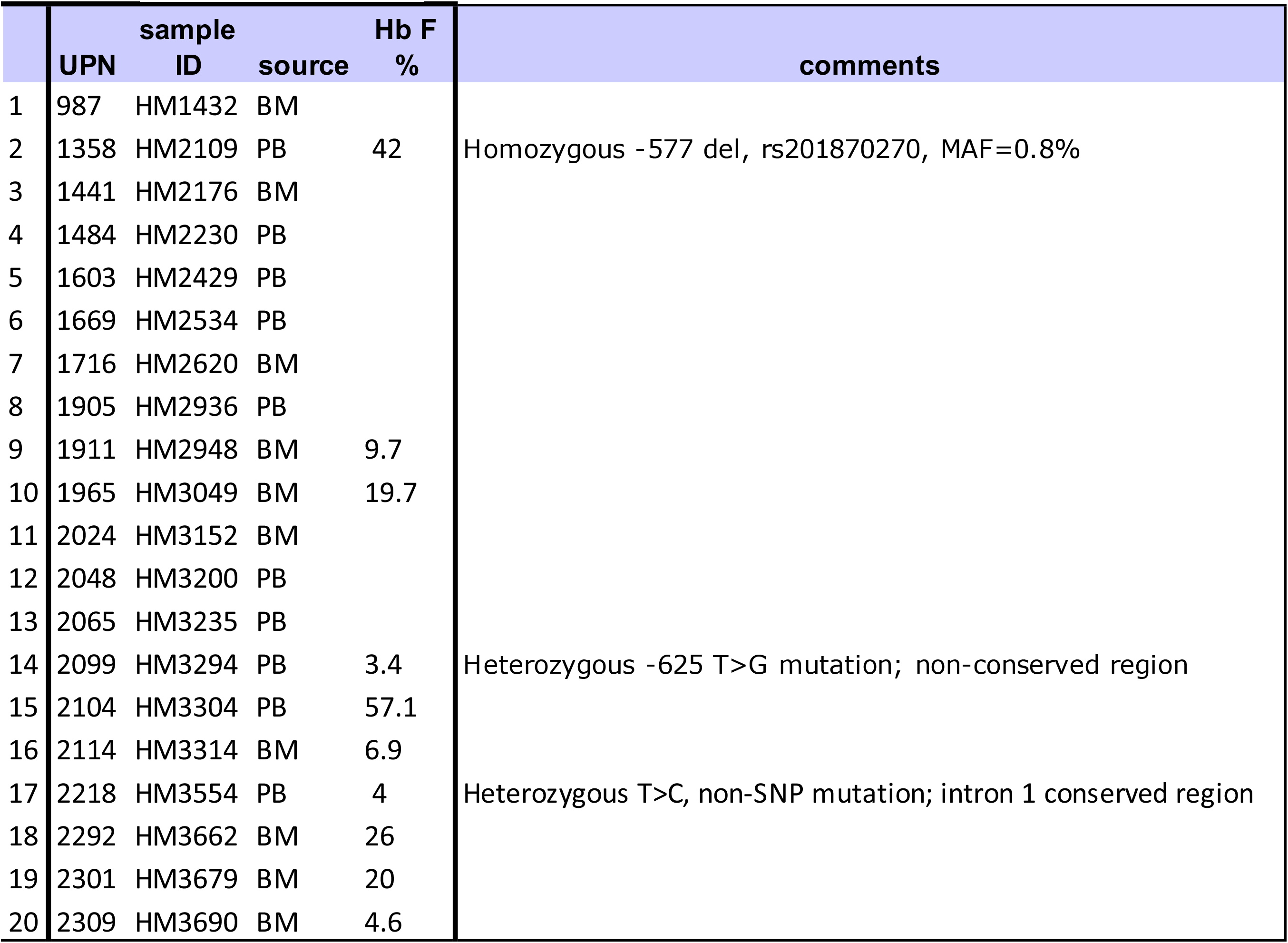
JMML patient samples.

**TABLE S2.**
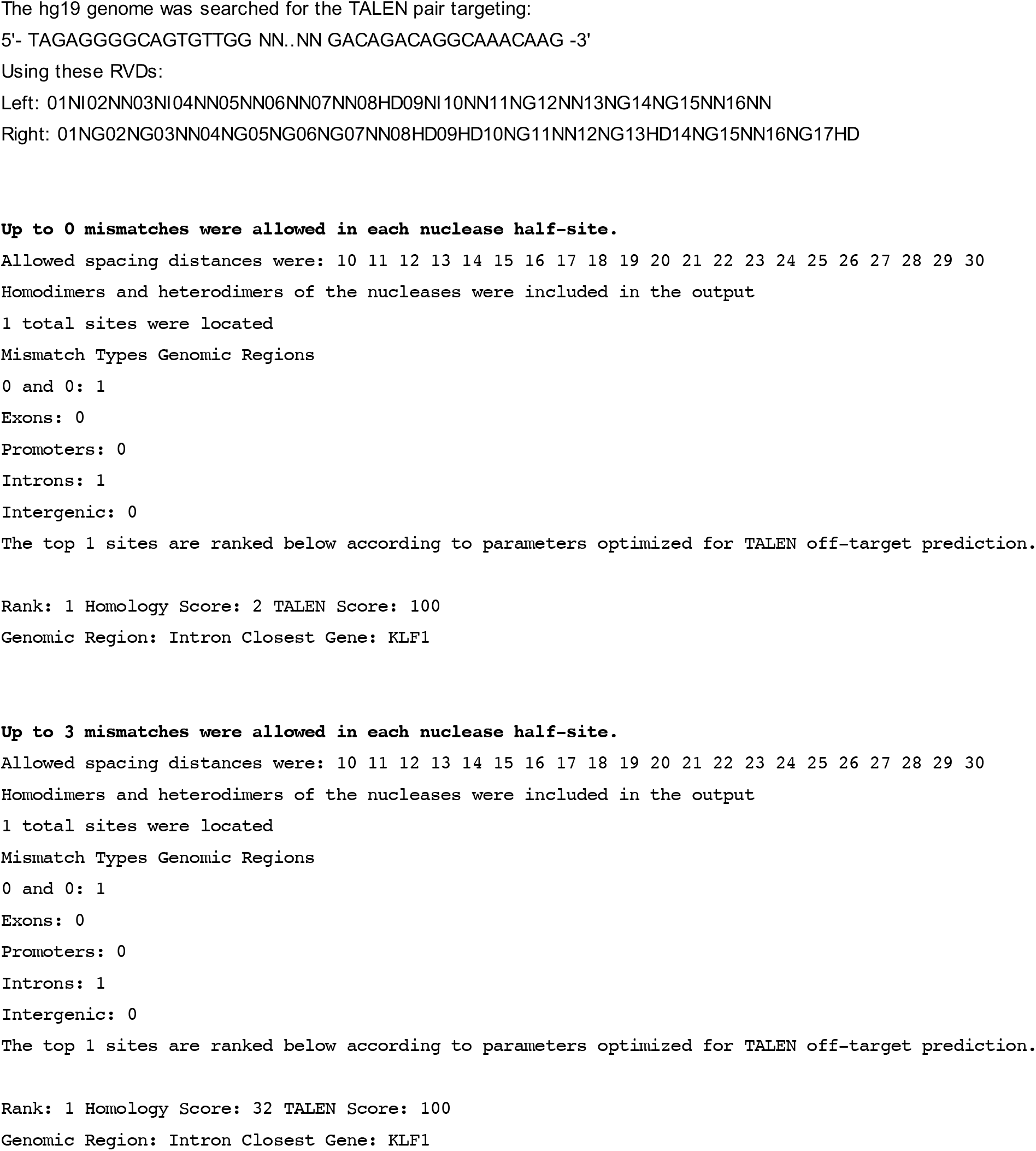
PROGNOS Summary for _home_apache_public_html_bao_Research_BioinformaticTools_prognos_03272019OHU99nls_Ranking-TALENv2.0.

**TABLE S3.**
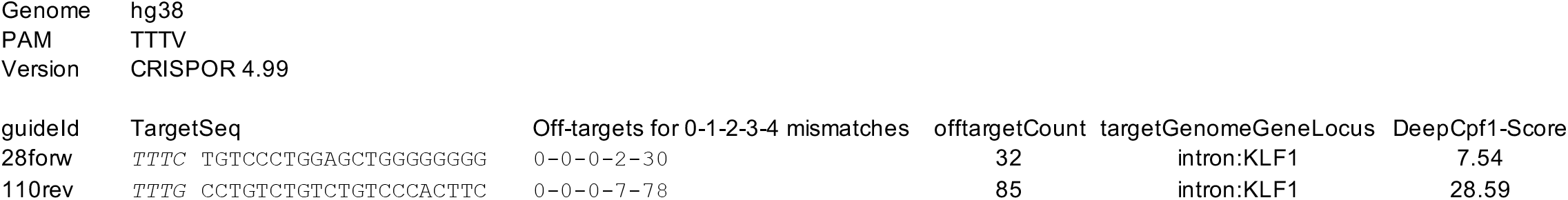
CRISPOR analysis of gRNA designs.

